# Transformation of the drug ibuprofen by *Priestia megaterium*: Reversible glycosylation and generation of hydroxylated metabolites

**DOI:** 10.1101/2024.03.18.585558

**Authors:** Tjorven Hinzke, Rabea Schlüter, Annett Mikolasch, Daniela Zühlke, Patrick Müller, Katharina Riedel, Michael Lalk, Dörte Becher, Halah Sheikhany, Frieder Schauer

## Abstract

As one of the most-consumed drugs worldwide, ibuprofen (IBU) reaches the environment in considerable amounts as environmental pollutant, necessitating studies of its further biotransformation as potential removal mechanism. Therefore, we screened bacteria with known capabilities to degrade aromatic environmental pollutants, belonging to the genera *Bacillus*, *Priestia* (formerly also *Bacillus*) *Paenibacillus*, *Mycobacterium*, and *Cupriavidus*, for their ability to transform ibuprofen. We identified five transformation products, namely 2-hydroxyibuprofen, carboxyibuprofen, ibuprofen pyranoside, 2-hydroxyibuprofen pyranoside, and 4-carboxy-α-methylbenzene-acetic acid. Based on our screening results, we focused on ibuprofen biotransformation by *Priestia megaterium* SBUG 518 with regard to structure of transformation products and bacterial physiology. Biotransformation reactions by P. megaterium involved (A) the hydroxylation of the isobutyl side chain at two positions, and (B) conjugate formation via esterification with a sugar molecule of the carboxylic group of ibuprofen and an ibuprofen hydroxylation product. Glycosylation seems to be a detoxification process, since the ibuprofen conjugate (ibuprofen pyranoside) was considerably less toxic than the parent compound to *P. megaterium* SBUG 518. Based on proteome profile changes and inhibition assays, cytochrome P450 systems are likely crucial for ibuprofen transformation in *P. megaterium* SBUG 518. The toxic effect of ibuprofen appears to be caused by interference of the drug with different physiological pathways, including especially sporulation, as well as amino acid and fatty acid metabolism.

**Importance:** Ibuprofen is a highly consumed drug, and, as it reaches the environment in high quantities, also an environmental pollutant. It is therefore of great interest how microorganisms transform this drug and react to it. Here, we screened several bacteria for their ability to transform ibuprofen. *Priestia megaterium* SBUG 518 emerged as highly capable and was therefore studied in greater detail. We show that *P. megaterium* transforms ibuprofen via two main pathways, hydrolyzation and reversible conjugation. These pathways bear resemblance to those in humans. Ibuprofen likely impacts the physiology of *P. megaterium* on several levels, including spore formation. Taken together, *P. megaterium* SBUG 518 is well suited as a model organism to study bacterial ibuprofen metabolism.

## Introduction

Pharmaceuticals, albeit intended as being beneficial to diagnose and treat illnesses, can not only have harmful side effects for humans, but can also be detrimental for ecosystems (1–5). Ibuprofen (IBU) as pain-mitigating and non-steroidal anti-inflammatory drug (NSAID) is the most commonly used analgesic in Germany (6) and the world’s third-most consumed drug (7). A main source of environmental contamination are wastewater treatment plant effluents: despite that IBU is removed from wastewater with comparatively high efficiency of around 90 % or more (1, 8–10), it is frequently detected in the effluents, as well as in surface waters (Weigel et al., 2004; Wojcieszyńska et al., 2022; Zuccato et al., 2005), and even in the Antarctic (13).

While IBU transformation and degradation in the environment are thus of great interest, microbial IBU transformation and degradation pathways are still largely unknown (14–17). Studies regarding IBU transformation by microorganisms include foremost whole-community degradation studies in bioreactors (18–20), and activated sludge communities (21–23). Some bacterial strains which degrade IBU are described, including *Patulibacter* sp. I11 (24), a *Nocardia* sp. (25), *Sphingomonas* sp. Ibu-2 (26), *Variovorax* sp. Ibu-1 (27), *Rhizorhabdus* (*Sphingomonas*) *wittichii* MPO218 (28, 29), *Sphigopyxis granuli* (30), *Rhodococcus cerastii* (31), *Bacillus thuringiensis* B1 (32), and *Rhizobium daejeonense* IBU_18 (33). Additionally, while IBU has antimicrobial effects (34–36), the impact of this drug on bacterial physiology and metabolism still needs to be elucidated.

To shed light on microbial IBU degradation pathways and molecular reactions to this drug in bacteria, we conducted a screening experiment to test several bacterial strains for their ability to transform IBU. A total of seven bacterial strains were chosen for testing based on known capability for environmental pollutant degradation. These strains belonged to the genera *Bacillus*, *Priestia*, *Paenibacillus*, *Mycobacterium* and *Cupriavidus*. Incubations with *Priestia megaterium* SBUG 518 (formerly *Bacillus megaterium* SBUG 518; (37)) yielded the most transformation products; therefore we performed further in-depth analyses with this strain. Our analyses encompassed elucidating IBU transformation pathways, as well as analysing the strains’ global proteomic responses to IBU.

We here show that *P. megaterium* SBUG 518 hydroxylates IBU, and, in addition, glycosylates not only IBU, but also at least one IBU oxidation product. We demonstrate that the direct glycosylation of IBU is reversible.

Furthermore, changes in the proteomic profile of *P. megaterium* SBUG 518 suggest that a *P. megaterium*-specific cytochrome P450 system is involved in IBU transformation, and that IBU interferes with sporulation, amino acid and fatty acid metabolism and potentially the oxidative stress response, providing a molecular basis for the toxic effect of IBU on *P. megaterium* SBUG 518.

## Material and Methods

### Strains

For screening experiments, we tested seven bacterial strains with known transformation capabilities for their effectiveness to transform IBU (Table 1). The strains are deposited into the strain collection of the Department of Biology of the University of Greifswald (SBUG).

**Table 1:** Bacterial strains used in this study, their origin and transformation capabilities of these or related strains. ATCC: American Type Culture Collection; DSM: Deutsche Sammlung von Mikroorganismen.

| Strain | Origin/ original strain number | Transformation capabilities |
| --- | --- | --- |
| *Priestia megaterium* SBUG 518 | ATCC 2007 | Oxidation of phenanthrene, fluoranthrene, pyrene (38) |
| *Priestia megaterium* SBUG 1979 | R. Biedendieck, Technical University Braunschweig, Institute of Microbiology, 01/2014 | Oxidation and hydroxylation of betulinic acid (39)<br>Dichloroanilines as C- and energy source (40) |
| *Bacillus pumilus* SBUG 1800 | isolated from Rassower Strom (41) | Vanillin degradation, including ring cleavage (42)<br>Ferulic acid decarboxylation (43) |
| *Bacillus thuringiensis* SBUG 1431 | DSM 350 | Dimethyl phthalate as C source (44) |
| *Cupriavidus basilensis* SBUG 290 | (45) | Dibenzofuran degradation (46)<br>Bisphenol transformation and ring cleavage (47, 48) |
| *Mycobacterium neoaurum* SBUG 109 | (49) | Transformation of ibuprofen and acetaminophen (50, 51)<br>Growth on pristane (52) |
| *Paenibacillus apiarius* SBUG 1947 | Isolated from Lake Balschasch (53) | Growth on dibenzofuran (54) |

### Culture conditions for screening experiments

Culture conditions were chosen according to the respective organisms’ physiology and expert knowledge (data not shown). We cultivated all strains of the genera *Bacillus*, *Priestia*, and *Paenibacillus* in 500-mL-flasks, containing 100 mL nutrient broth (NB II) for 24 h at 30 °C and 180 rpm on a rotary shaker (HT FORS, Infors AG, Bottmingen, Switzerland). Both strains of *P. megaterium* were cultivated additionally in Lysogeny broth (LB).

Cultivation of *C. basilensis* with biphenyl was carried out following Zühlke et al., 2017, with the minor modification that the main culture was inoculated with only 5 ml cell suspension in 100 ml NB II.

*M. neoaurum* was cultivated on agar plates with tetradecane for 4 d at 30 °C, using material of five plates for biotransformation experiments as described by Nhi-Cong et al., 2009. Cells were transferred into 75 ml mineral salts medium for bacteria (MMb; pH 6.3; according to Hundt et al., 1998) supplemented with IBU (see below).

### Culture conditions for experiments with *P. megaterium* SBUG 518

Cultivation for incubation experiments with stationary-phase cells: We grew bacteria on nutrient agar for 24 h at 30 °C and then transferred the cells into 100 mL NB II or LB in 500-mL-flasks. Microorganisms were cultivated for 24 h at 30 °C and 180 rpm on a rotary shaker.

Cultivation for incubation experiments with logarithmic-phase cells: Bacteria were grown on nutrient agar for 12 h at 30 °C. Subsequently, we transferred the cells into 100 mL NB II in 500-mL-flasks. The cells were cultivated for 5 h at 30 °C and 180 rpm on a rotary shaker until an optical density OD_500nm_ of 1.0 to 1.5 was reached.

### Biotransformation experiments

Transformation experiments with IBU were performed in 500-mL-flasks containing 100 mL MMb and 0.005% [0.05 mg mL^-1^; equivalent to 219 μM] IBU. For incubation experiments in the presence of glucose, we carried out transformation experiments in 500-ml-flasks containing 100 ml MMb, 0.005% IBU and 0.1% glucose (added from a 50% aqueous glucose stock solution [w/v]).

After cultivation (see above), we harvested the cells by centrifugation (11,270 x g, 10 min, 4 °C), washed them twice with MMb and resuspended them in a small amount of MMb. We then added this cell suspension to the flasks with MMb and IBU to reach an optical density at 500 nm (OD_500nm_) of 3.0, corresponding in the case of *P. megaterium* SBUG 518 to a cell titre of approximately 3.95*10^8^ [±0.55*10^8^] cells mL^-1^. Incubation experiments were carried out on a rotary shaker at 30 °C and 130 rpm (VKS-75 Control, Edmund Bühler GmbH, Bodelshausen, Germany).

Two types of controls were used: (i) flasks with cells in MMb without IBU and (ii) flasks with IBU in MMb without cells.

### Cytochrome P450 inhibition studies

To study the effect of cytochrome P450 inhibition on IBU biotransformation by *P. megaterium* SBUG 518, we added 219 μM 1-aminobenzotriazole to 500-ml-flasks filled with 100-ml MMb containing 219 μM IBU (0.005%) in the presence and absence of 0.1% glucose. After inoculation with stationary-phase cells (see section “Biotransformation experiments”) flasks were shaken on a rotary shaker at 30 °C and 130 rpm.

### Analysis of IBU biotransformation with high-performance liquid chromatography (HPLC)

To study IBU biotransformation, including the formation of transformation products, over the time course of the incubations, we used 1-ml samples of culture supernatant taken under sterile conditions at selected time points. We removed cells via centrifugation (3,600 x g, 10 min, Hettich Universal 30F, Tuttlingen) and analyzed 60 µl of the resulting supernatant by HPLC using an Agilent-Technologies 1200 Series system (Santa Clara, USA). Products were separated on a LiChroCART 125-4 RP-18 end-capped 5 µm column (Merck, Darmstadt, Germany) with a solvent system of methanol and phosphoric acid (0.1%, v/v) with a linear gradient from 30% to 100% methanol over a period of 14 min at a flow rate of 1 mL min^-1^.

### Extraction of biotransformation products

For purification of transformation products, we harvested culture supernatants by centrifugation (3,600 x g, 5 min, Hettich Universal 30F) and extracted the supernatants four times with twice the volume of ethyl acetate at pH 7 and again four times with twice the volume of ethyl acetate at pH 2. A 25% (w/v) aqueous sodium hydroxide solution was used to adjust the pH value to pH 7; the aqueous supernatant was then adjusted to pH 2 with 32% (v/v) hydrochloric acid. Organic phases were dried with anhydrous sodium sulfate. After rotary evaporation, we resolved the residues in methanol and stored them at −20 °C until further analysis.

### Chemical analysis, isolation, and identification of IBU transformation products

IBU transformation products were purified on an Agilent Technologies 1260 Infinity semi-preparative HPLC (Santa Clara, USA) with an Eclipse XDB-C18, PN 977 250-102, 21.2 x 250 mm; 7 mm column (Agilent, Santa Clara, USA), using an 8 min linear gradient of 40% to 100% methanol in acetic acid (0.1% v/v) at a flow rate of 10 mL min^-1^. The isolated products were concentrated by rotary evaporation, the residues dissolved in methanol and stored at −20 °C until further analysis.

An Agilent Technologies 1200 Series 6120 Quadrupole model was used for liquid chromatography-mass spectrometry (LC-MS) analysis. A ZORBAX SB-C18 column (2.1 x 50 mm, pore size 1.8 mm) was used for HPLC separation at a flow rate of 0.1 mL min^-1^ with a 7 min gradient from 10% to 100% acetonitrile in 0.1% aqueous ammonium formate. The MS was used with an electrospray ionization (API-ES) source (dry and nebulizer gas: nitrogen; drying gas flow 10.0 l min^-1^; nebulizer pressure 45 psig; drying gas temperature 350 °C; capillary voltage 4000 V).

We used the methods described by Mikolasch et al. (2016) for the analyses of extracts by gas chromatography coupled to mass spectrometry (GC-MS). In brief, IBU biotransformation products were detected by injecting 1 µL of the extracts of the biotransformation experiments into an Agilent gas chromatograph 7890A GC System (Waldbronn, Germany) equipped with a capillary column (Agilent 1901 S-433, 30 m x250 µm x 0.25 µm, HP-5ms column) and a mass selective detector 5975C inert XL EI/CI MSD with a quadrupole mass spectrometer.

The nuclear magnetic resonance (NMR) spectra of two products (P3, P4; see Results) dissolved in dimethyl sulfoxide (DMSO)-d6 were obtained on a Bruker Avance-II instrument (Bruker Biospin GmbH, Rheinstetten, Germany) at 600 MHz (^1^H, ^13^C, heteronuclear multiple bond correlation (HMBC), heteronuclear single quantum coherence (HSQC)).

### Toxicity studies

To estimate the toxicity of IBU and IBU pyranoside (IBU-PYR, product P3) on *P. megaterium* SBUG 518, we performed growth assays with and without these substances in NB II (pH 6.0; adjusted with HCl from an initial pH of 7.2). We decreased the pH to allow for greater IBU toxicity effects against *P. megaterium* SBUG 518. Cells were cultivated in 500-mL-flasks containing 100 mL NB II for 24 h at 30 °C and 180 rpm on a rotary shaker. The cell suspension was transferred into 100-mL-Erlenmeyer-flasks containing 20 mL NB II, pH 6.0 (OD_500nm_ = 0.2), and 43.8 µM IBU (corresponding to a growth inhibitory concentration of 0.05% IBU) or 43.8 µM of product P3, respectively, were added. Controls without IBU or IBU transformation product were included in the assays. Subsequently, we cultivated the assays for 72 h at 30 °C and 180 rpm, and periodically determined the OD_500nm_.

### Analysis of impacts of IBU on the proteome of *P. megaterium* SBUG 518

To analyse changes in the *P. megaterium* SBUG 518 proteome profile after incubation with IBU, we took 10-mL-samples of *P. megaterium* SBUG 528 IBU transformation assays conducted with stationary-phase cells and of control incubation without IBU after 1 h and 24 h of incubation. We immediately placed samples on ice, harvested cells by centrifugation (10,015 x g, 5 min, 4 °C, Heraeus Biofuge Primo R, Thermo Scientific), washed the cell pellets twice in TE buffer (10 mM Tris, 2 mM EDTA, pH 7.5) and subsequently resuspended the cells in TE buffer with 1% (w/v) Triton-X-100. For cell lysis, 500 µl glass beads (diameter 0.10–0.11 mm, Sartorius AG, Göttingen, Germany) were added to 1 mL sample and cells disrupted with a FastPrep®24 homogenizer (M. P. Biomedicals, Irvine, California, USA) in three 30 s cycles at 6.5 m s^-1^ with cooling samples on ice for 5 min between cycles. To remove glass beads and cell debris, samples were centrifuged twice (5 min, 21,885 x *g*, 4 °C, Heraeus Biofuge Primo R) and the supernatant transferred to a new tube. Proteins were precipitated overnight with six times the sample volume of ice-cold acetone at −20 °C. Subsequently, we pelleted proteins by centrifugation for 1 h at 10,015 x g and 4 °C, discarded the supernatant and resuspended the protein pellet in 1 mL ice-cold acetone. After repeating the washing step, we dried the protein pellets at room temperature and then resuspended them in 0.5 mL 8 M urea/ 2 M thiourea. Following centrifugation for 10 min at 10,015 x g and room temperature, the supernatant was transferred into a new tube. Protein concentration was determined with Roti^®^-Nanoquant (Roth, Karlsruhe, Germany) reagent according to the manufacturer’s instructions. Protein solutions were stored at −20 °C. Experiments were performed in triplicate. Proteins were separated using 1D-SDS-PAGE as described in Zühlke et al., 2017, using precast 4-20 % polyacrylamide gels (4-20 % Criterion^TM^ TGX BioRad^TM^, BIO-RAD, Hercules, USA).

Liquid chromatography-tandem mass spectrometry (LC-MS-MS) analysis was performed using a nanoACQUITY^TM^-UPLC^TM^-System (Waters, Milford, USA) combined with a linear trap quadrupole (LTQ)-Orbitrap mass spectrometer (Thermo Fisher Scientific, Waltham, USA). Peptides were loaded on a pre-column (Symmetry C18, 5 μm, 180 μm inner diameter x 20 mm, Waters, Milford, USA) and washed for 3 min at a flow rate of 10 μL min^-1^ with 0.1 % acetic acid. Peptides were then eluted onto the analytical column (BEH130 C18, 1.7 μm, 100 μm inner diameter x 100 mm, Waters, Milford, USA) with a step-wise gradient of 5% to 99% acetonitrile in 0.1% acetic acid in 80 min at a flow rate of 400 nL min^-1^ (0.5% acetonitrile min^-1^ to 25% acetonitrile, 0.83% acetonitrile min^-1^ to 50% acetonitrile, 49% acetonitrile min^-1^ to 99% acetonitrile in 0.1% acetic acid). Measurements were performed in LTQ/Orbitrap parallel mode. Survey scans were performed in the orbitrap (m/z range 300–2000, resolution 30,000, lock mass option enabled, lock mass 445.120025). Per scan, the five peaks with highest intensity were fragmented in the LTQ using collision induced dissociation (CID). Precursor ions with unknown as well as those with single charge were excluded from fragmentation. Dynamic exclusion for precursor ions was set to 30 s.

To identify detected proteins, raw spectra were searched against a forward-reverse database of *P. megaterium* ATCC 14581 (20.11.2017, Uniprot) with added common laboratory contaminants. Sorcerer^TM^-SEQUEST^®^ and Scaffold_4 were used, with the following parameters: trypsin (KR), maximum two omitted cleavage sites, precursor mass monoisotopic, precursor mass range 400–4.500 Da, 1 Da fragment mass accuracy, b- and y-ion series, peptide mass tolerance 10 ppm, variable oxidation of methionine (15.99 Da) with a maximum of four modifications per peptide. The search result was filtered with “XCorr versus charge state”-filters as follows: 2.2 for twofold charged, 3.3 for threefold and 3.75 for fourfold and higher charged ions and DeltaCn 0.1.

#### Pride upload references

The mass spectrometry proteomics data have been deposited to the ProteomeXchange Consortium via the PRIDE (58) partner repository with the dataset identifier PXD015716 (access for reviewers: https://www.ebi.ac.uk/pride/archive/login, username, password: 0yxboXRw)

#### Statistical evaluation and functional annotation of proteomics results

Proteins were quantified by calculating Normalized Spectral Abundance Factors (NSAFs) (59). Statistical analysis was performed in R (60) using the package limma v. 3.50.3 (61). Correction of p-values for multiple comparisons was done using Benjamini-Hochberg correction (α=0.05) (62) implemented in limma. Additionally, we required proteins to have at least a 2-fold abundance change in the IBU incubation vs. control at the same sampling timepoint to be considered significantly changed. For visualization, we employed the package EnhancedVolcano 1.12.0 (63).

### Chemicals

IBU sodium salt (molecular weight of 228.26 g mol^-1^), 2-hydroxyibuprofen (2-OH-IBU) and carboxyibuprofen (CBX-IBU) were obtained from Sigma-Aldrich (Steinheim, Germany). All chemicals and solvents used were of the highest purity available.

## Results

### Screening of bacterial strains

All of the seven screened bacterial strains transformed IBU, as determined by HPLC analysis (Table 2). Five different transformation products, P1 to P5, were identified based on HPLC and LC-MS analyses. All of the organisms produced P1, while none produced all five transformation products.

**Table 2:**
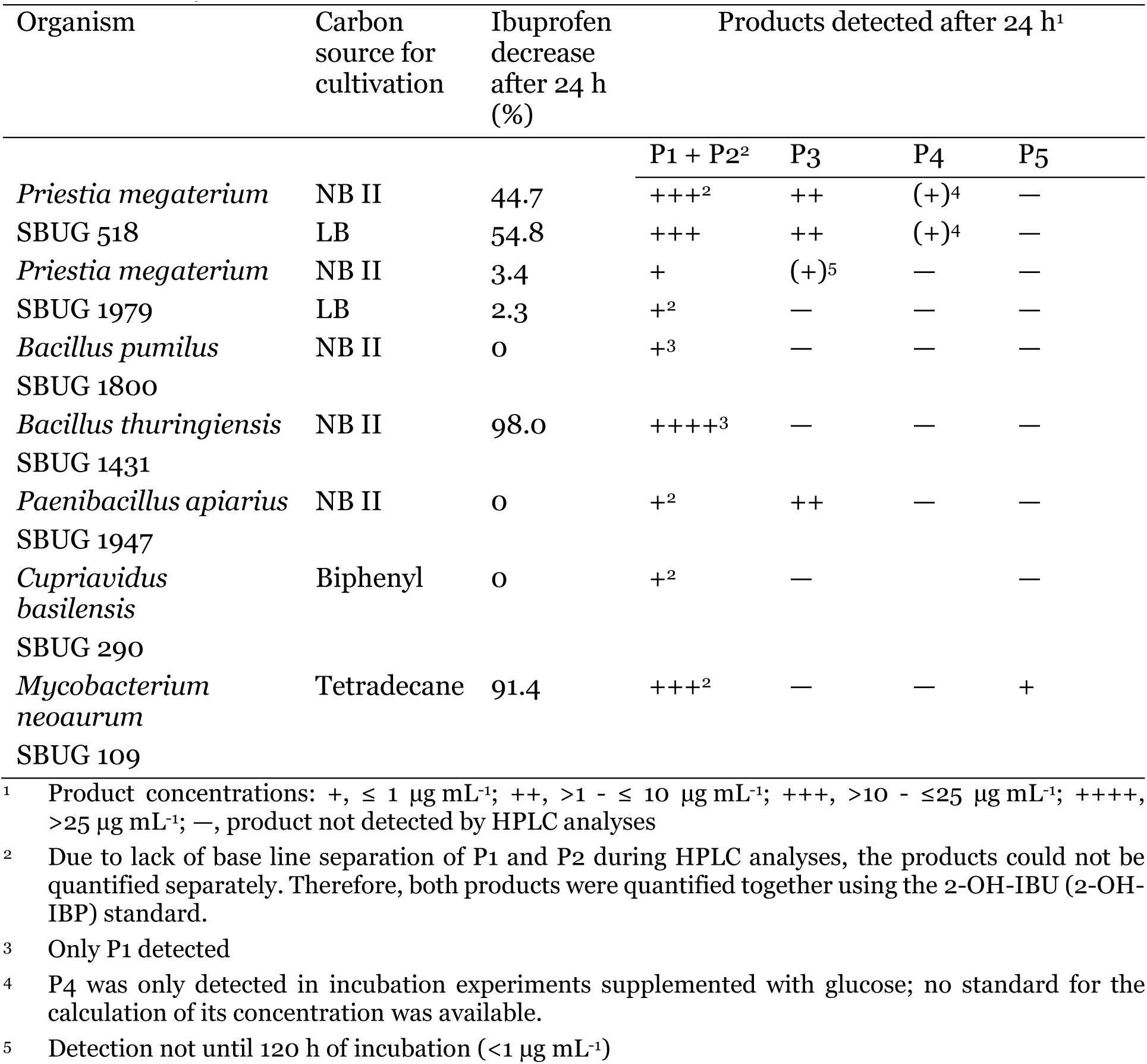
Biotransformation of ibuprofen and transformation products formed during the transformation of 0.005% ibuprofen by different bacterial strains. Shown are the relative decrease of IBU concentration and detected transformation products in the culture supernatant after 24 h of incubation of various bacterial strains with 0.005% IBU, cultivated with different carbon sources prior to incubation.

*P. megaterium* SBUG 518 produced the highest number and a high yield of transformation products. Therefore, we chose *P. megaterium* SBUG 518 as model organism for further studies.

### Structure elucidation of transformation products

Products formed during the incubation of the bacterial strains with IBU were identified by HPLC analyses via comparison of the UV-VIS spectrum and retention time with the data of authentic standards, as well as by GC-MS, LC-MS (Supp. Table S1) and/or NMR analyses (Table 3, Supp. Tables S2 and S3).

**Table 3:**
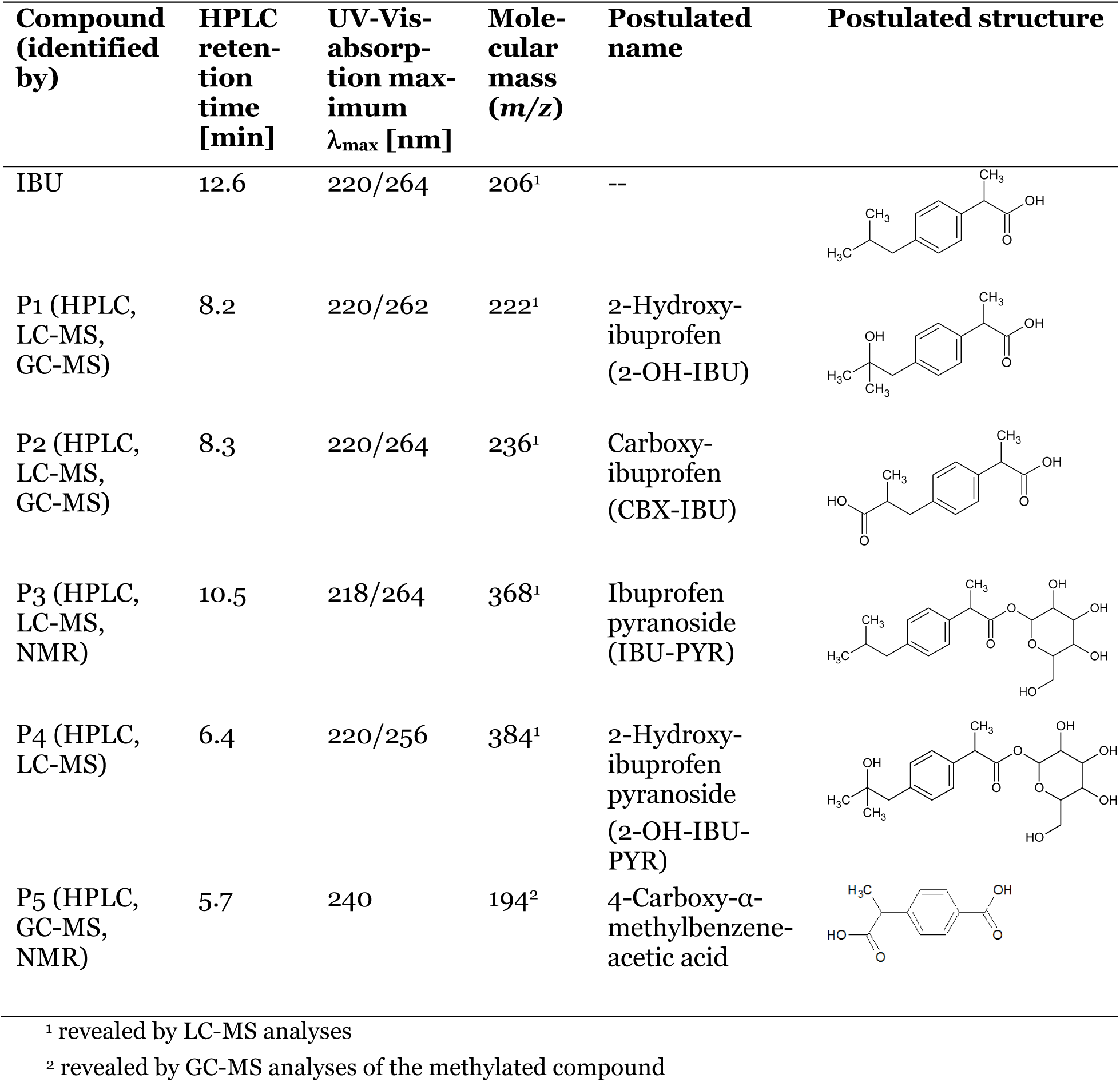
Transformation products formed during the transformation of IBU by various bacteria, identified by HPLC, LC-MS, GC-MS, and NMR analyses. Shown here are HPLC analysis results. For GC-MS and LC-MS results, see Supp. Table S1, for NMR results, see Supp. Tables S2 and S3.

| Compound (identified by) | HPLC retention time [min] | UV-Vis-absorption maximum $\lambda_{\max}$ [nm] | Molecular mass (m/z) | Postulated name | Postulated structure |
| --- | --- | --- | --- | --- | --- |
| IBU | 12.6 | 220/264 | 206<sup>1</sup> | -- |  |
| P1 (HPLC, LC-MS, GC-MS) | 8.2 | 220/262 | 222<sup>1</sup> | 2-Hydroxy-ibuprofen (2-OH-IBU) |  |
| P2 (HPLC, LC-MS, GC-MS) | 8.3 | 220/264 | 236<sup>1</sup> | Carboxy-ibuprofen (CBX-IBU) |  |
| P3 (HPLC, LC-MS, NMR) | 10.5 | 218/264 | 368<sup>1</sup> | Ibuprofen pyranoside (IBU-PYR) |  |
| P4 (HPLC, LC-MS) | 6.4 | 220/256 | 384<sup>1</sup> | 2-Hydroxy-ibuprofen pyranoside (2-OH-IBU-PYR) |  |
| P5 (HPLC, GC-MS, NMR) | 5.7 | 240 | 194<sup>2</sup> | 4-Carboxy-α-methylbenzeneacetic acid | |

<sup>1</sup> revealed by LC-MS analyses
<sup>2</sup> revealed by GC-MS analyses of the methylated compound

### Biotransformation experiments with *P. megaterium* SBUG 518

#### Biotransformation of IBU in the absence of glucose leads to three main transformation products

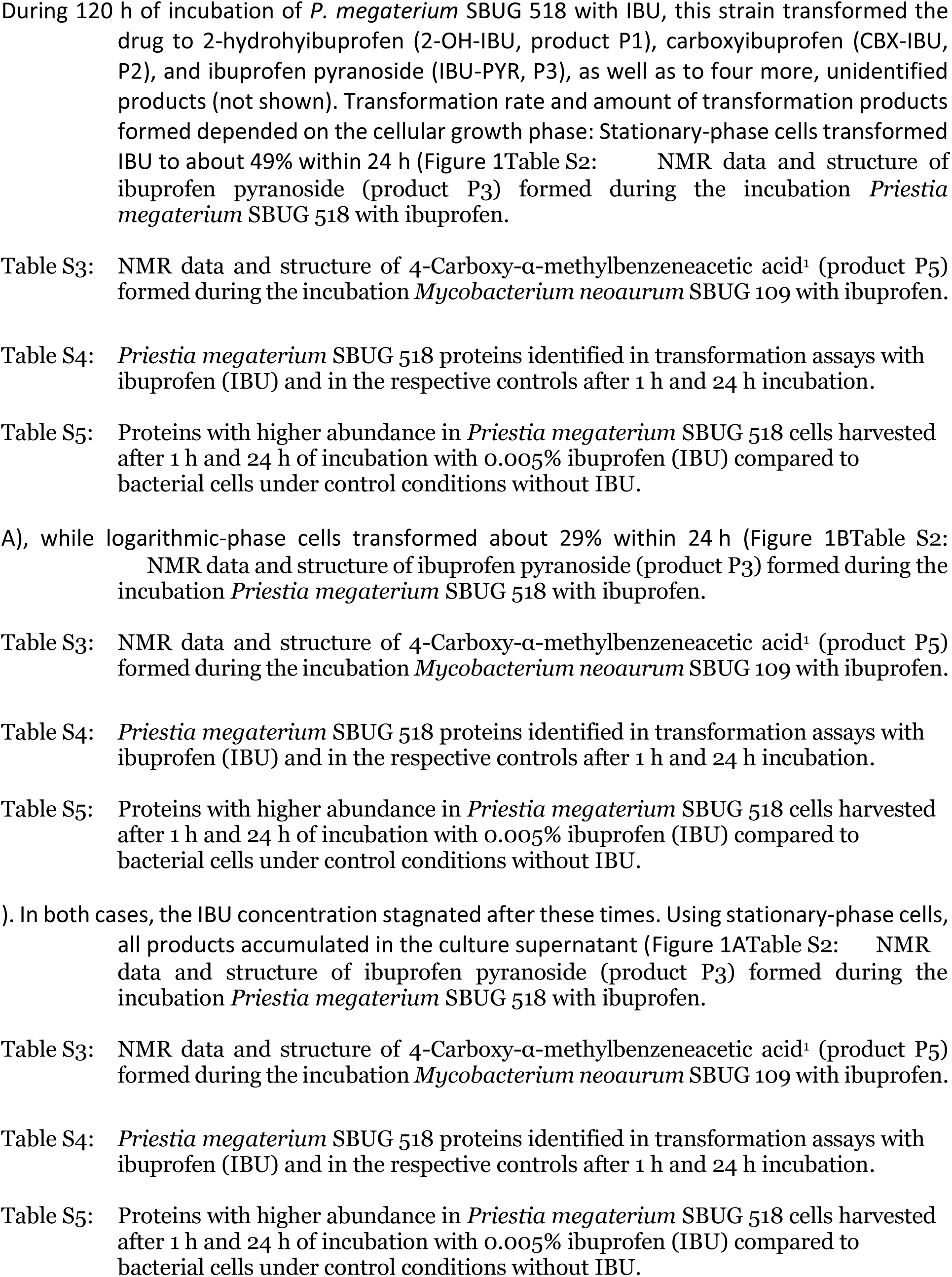
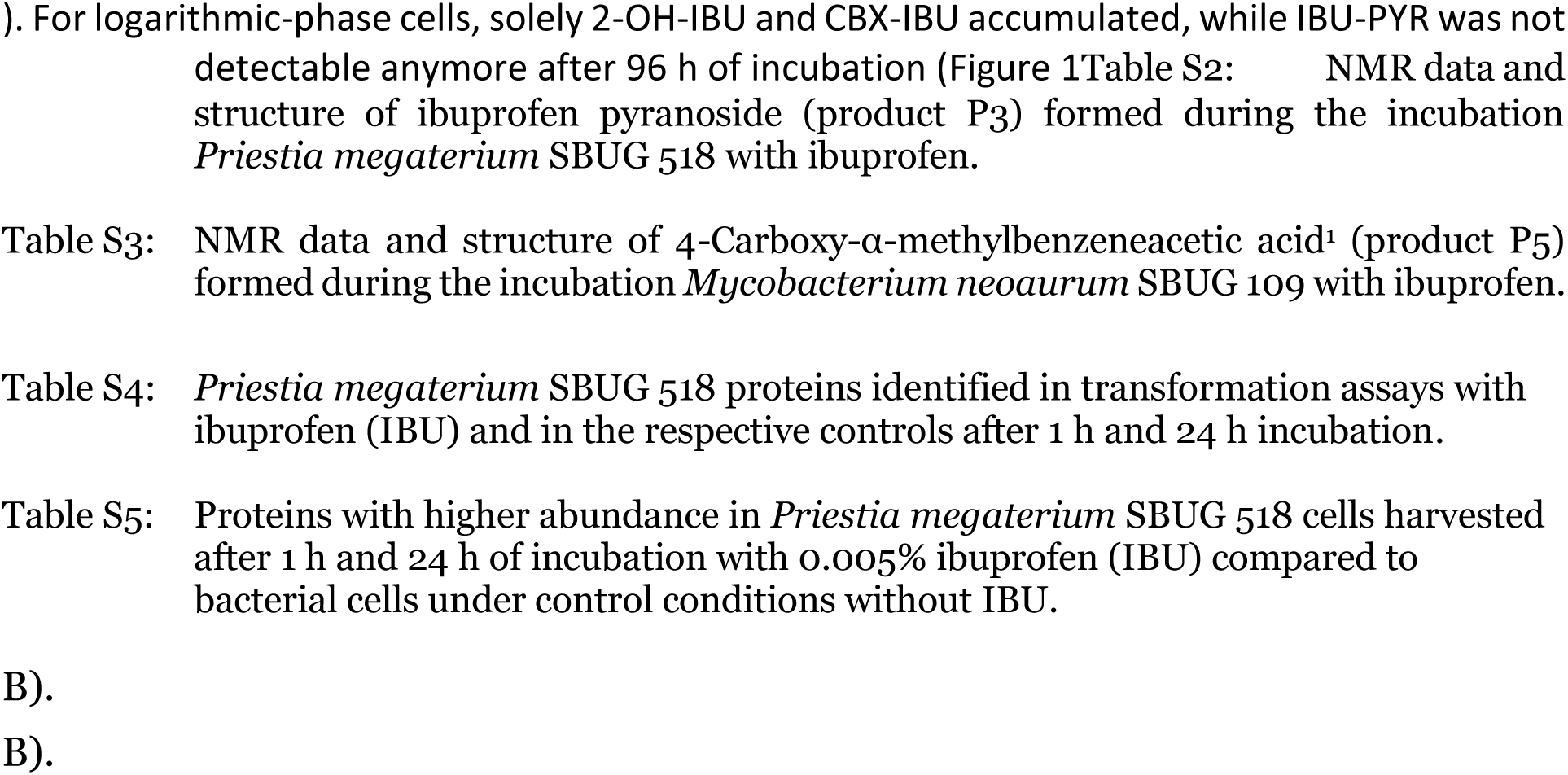

#### Biotransformation of IBU in the presence of glucose yields higher concentrations of IBU-PYR

After incubation of stationary-phase cells of *P. megaterium* SBUG 518 with IBU in the presence of 0.1% glucose, we detected the transformation product 2-OH-IBU-PYR (product P4) by HPLC analysis, in addition to 2-OH-IBU, CBX-IBU, and IBU-PYR. Stationary-phase cells transformed about 92 % of IBU within the first 4 h of incubation, which was correlated with a sharp concentration increase of IBU-PYR (Figure 1C). After 4 h, IBU concentration increased again, whereas that of IBU-PYR decreased (Figure 1C). Thus, while about 49% of IBU was glycosylated after 4 h, after 24 h only about four percent of the drug remained glycosylated. The concentration of 2-OH-IBU-PYR increased in the same fashion as the IBU-PYR concentration, but with a 2 h-delay, and only slightly decreased in concentration after 6 h incubation (Figure 2).

**Figure 1:**
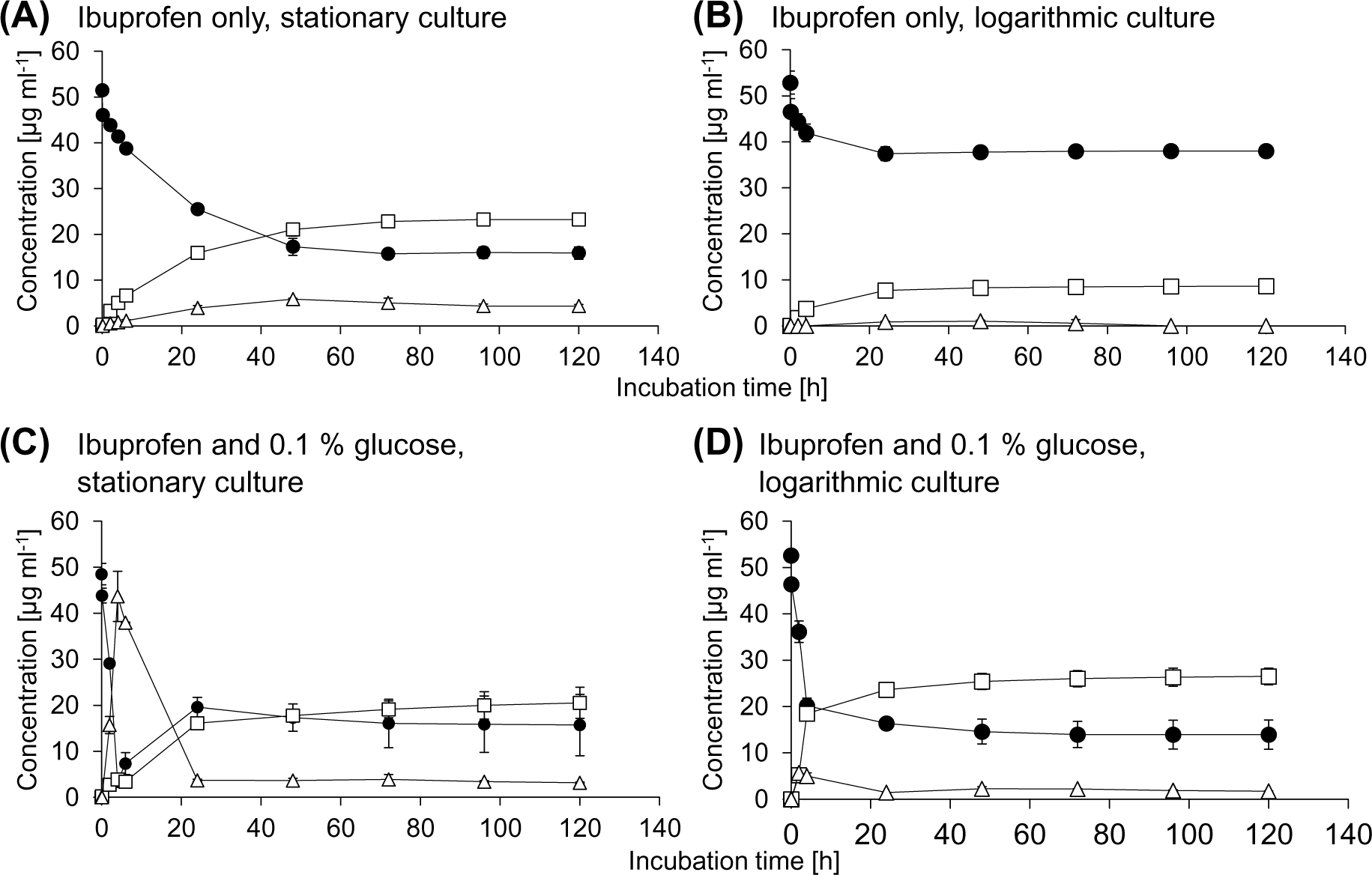
Time course of the biotransformation of 0.005 % IBU (black circles) by *Priestia megaterium* SBUG 518 and formation of the transformation products 2-OH-IBU and CBX-IBU (squares), and IBU-PYR (triangles). Incubation was carried out with (A) stationary-phase cells, and (B) logarithmic-phase cells in the absence of glucose as well as with (C) stationary-phase cells, and (D) logarithmic-phase cells in the presence of 0.1% glucose. Means and standard deviations of two independent parallels are shown.

**Figure 2:**
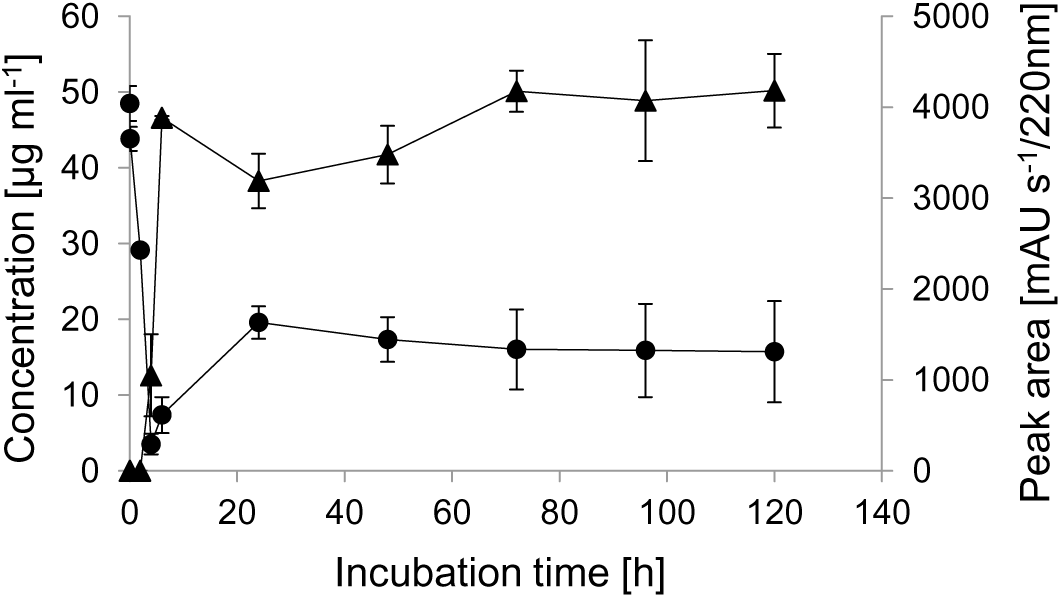
Time course of the biotransformation of IBU (circles, primary axis) and the formation of 2-OH-IBU-PYR (triangles, secondary axis) by stationary-phase cells of *Priestia megaterium* SBUG 518. Means and standard deviations of two independent parallels are shown.

Logarithmic-phase cells transformed about 73% IBU within 72 h to 2-OH-IBU, CBX-IBU, and IBU-PYR, with the highest transformation rate within the first 4 h of incubation (Figure 1D). All products accumulated in the culture supernatant until the end of incubation. The product 2-OH-IBU-PYR was detected only in one replicate after 4 h of incubation with logarithmic-phase cells.

### Cell cycle phase and glucose presence impact IBP biotransformation

When no glucose was present in the medium, stationary-phase and logarithmic-phase cells transformed about the same amount of IBU within the first 4 h of incubation. However, at the end of the incubation, stationary-phase cells had transformed almost three times more IBU and formed triple the amount of 2-OH-IBU and CBX-IBU, as well as fourfold more IBU-PYR than logarithmic-phase cells.

Glucose presence increased transformation rates, especially in the first 4 h. Stationary-phase cells produced more IBU-PYR than logarithmic-phase cells, but glycosylation was reversible only in incubations with stationary-phase cells. After 24 h incubation with IBU and glucose, product quantities of cells in both growth phases were similar. Please refer to Table 4 for a summary of IBU biotransformation results. In addition, IBU apparently impacted sporulation of *P. megaterium* SBUG 518: logarithmic-phase cells with IBU (but without glucose) produced only few spores after 120 h incubation, whereas under all other three conditions mostly spores where present after 120 h incubation (Supp. Figure S1).

**Table 4:** Concentration of ibuprofen (IBU) and of products formed after 4 h, 24 h, and 120 h incubation with IBU of stationary-phase cells and logarithmic-phase cells, respectively, in the absence and presence of 0.1% glucose in the incubation medium. Conc.: concentration; P1: 2-hydroxyibuprofen; P2: carboxyibuprofen, P3: ibuprofen pyranoside, P4: 2-hydroxyibuprofen pyranoside.

|  |  | IBU and transformation products formed by |  |  |  |  |  |  |  |  |  |
| --- | --- | --- | --- | --- | --- | --- | --- | --- | --- | --- | --- |
| incubation conditions | incubation time | Stationary-phase cells |  |  |  |  | Logarithmic-phase cells |  |  |  |  |
|  |  | IBU |  | P1+P2 | P3 | P4 | IBU |  | P1+P2 | P3 | P4 |
|  |  | Decrease of (%) | Conc. (mg mL<sup>-1</sup>) | Conc. (mg mL<sup>-1</sup>) |  | Peak area (mAU s<sup>-1</sup>/220 nm) | Decrease of (%) | Conc. (mg mL<sup>-1</sup>) | Conc. (mg mL<sup>-1</sup>) |  | Peak area (mAU s<sup>-1</sup>/220 nm) |
| Absence of glucose | 4 h | 10.2 | 41.4 | 5.0 | 0.8 | 0 | 9.8 | 41.9 | 3.8 | 0 | 0 |
|  | 24 h | 44.7 | 25.5 | 15.9 | 3.9 | 0 | 19.6 | 37.4 | 7.7 | 0.9 | 0 |
|  | 120 h | 65.4 | 15.9 | 23.2 | 4.3 | 0 | 18.2 | 38.0 | 8.6 | 0 | 0 |
| Presence of 0.1% glucose | 4 h | 91.9 | 3.5 | 3.9 | 43.7 | 1050 | 56.6 | 20.2 | 18.5 | 5.0 | 0 |
|  | 24 h | 55.4 | 19.6 | 16.1 | 3.8 | 3187.5 | 64.8 | 16.3 | 23.6 | 1.4 | 105.8 |
|  | 120 h | 64.4 | 15.7 | 20.5 | 3.2 | 4181.7 | 70.0 | 13.9 | 26.5 | 1.7 | 0 |

### 2-OH-IBU is glycosylated in the presence of glucose

When using 2-OH-IBU as biotransformation substrate in presence of glucose, stationary-phase cells formed 2-OH-IBU-PYR (product P4) with a very low transformation rate. Only about 7% of 2-OH-IBU was transformed by the cells within the first 6 h, during which time product concentration increased. Thereafter, product concentration decreased and substrate concentration increased (Figure 3).

**Figure 3:**
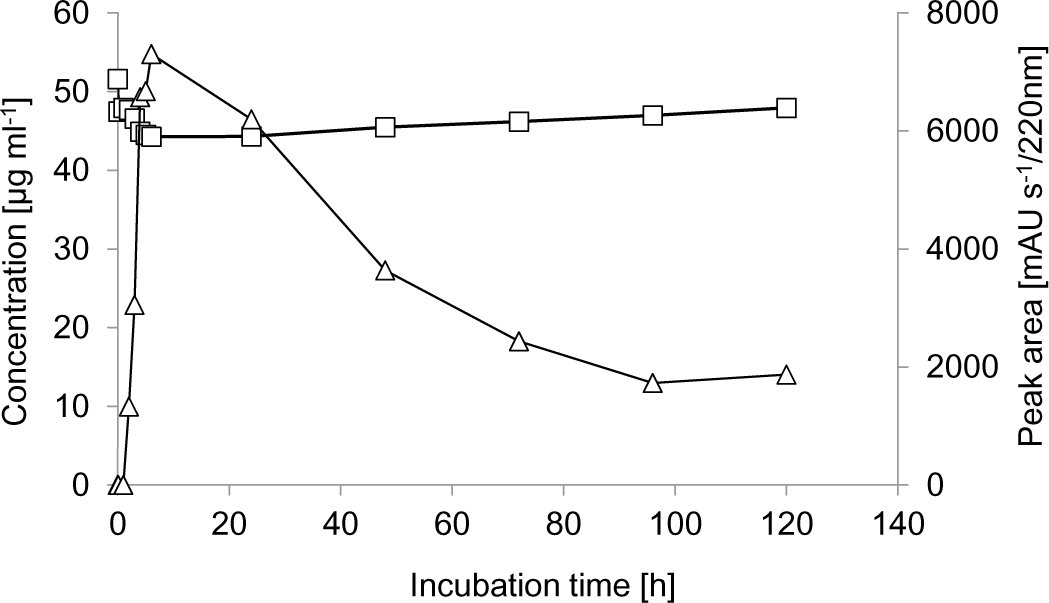
Time course of the biotransformation of 2-OH-IBU (squares) and the formation of 2-hydroxyibuprofen pyranoside (triangles) by *Priestia megaterium* SBUG 518. This experiment was carried out once.

### The transformation product IBU-PYR is less toxic than the drug IBU

IBU inhibited the growth of *P. megaterium* SBUG 518 at a concentration of 0.05% (equivalent to 2.19 mM) in NB II, pH 6.0 (but not in NB II with un-adjusted pH of 7.2, data not shown). However, with the transformation product IBU-PYR (P3) in equimolar concentration, growth of *P. megaterium* SBUG 518 was only slightly delayed compared to the control (Figure 4).

**Figure 4:**
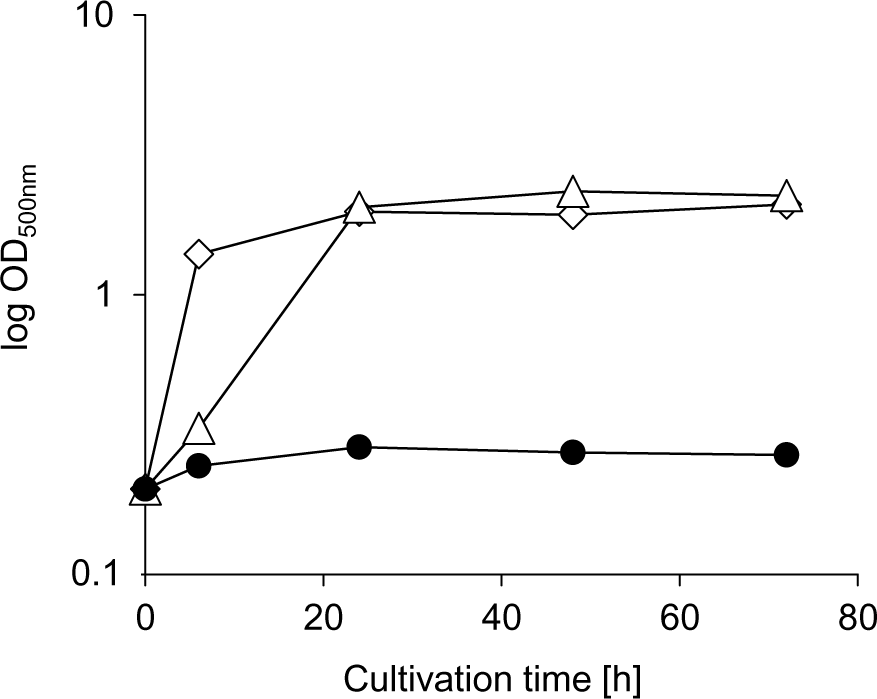
Effect of 0.05% (2.19 mM) IBU (black circles) and the equimolar concentration of the transformation product 2-hydroxyibuprofen pyranoside (triangles) on the growth of *Priestia megaterium* SBUG 518 in NBII (pH 6.0) compared to the control (diamonds) without IBU or transformation product. Shown are means and standard deviations of two independent parallels.

### Cytochrome P450 inhibition leads to decreased IBU oxidation

In the presence of the cytochrome P450 inhibitor 1-aminobenzotriazole, CBX-IBU and only about a tenth of the concentration of 2-OH-IBU as compared to the control without inhibitor was formed (Figure 5A). In contrast, the inhibitor had no effect on IBU-PYR concentrations.

**Figure 5:**
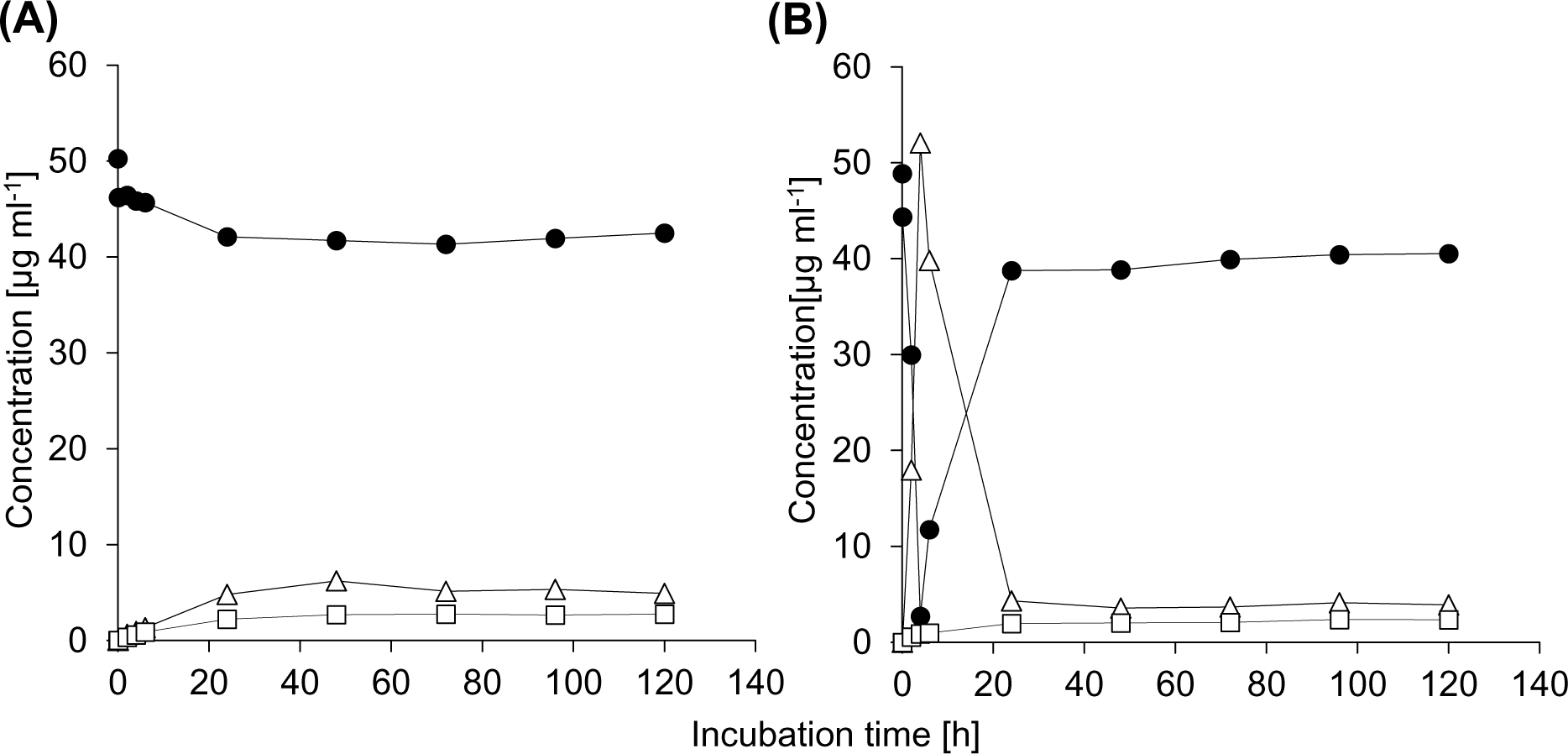
Time course of the biotransformation of IBU (black circles) by *Priestia megaterium* SBUG 518 and formation of the transformation products 2-OH-IBU (squares) and IBU-PYR (triangles) in the presence of (a) 219 μM 1-aminobenzotriazole and of (B) 219 μM 1-aminobenzotriazole and 0.1% glucose. These experiments were performed once.

Almost the same results were obtained in the presence of 1-aminobenzotriazole and 0.1% glucose. Only about one tenth of the concentration of 2-OH-IBU compared to control conditions, and neither CBX-IBU nor 2-OH-IBU-PYR were detected, whereas reversible formation of IBU-PYR occurred (Figure 5B).

### Influence of IBU on the proteome profile of *P. megaterium* SBUG 518

In total, we identified 1,346 *P. megaterium* SBUG 518 proteins after 1 and 24 h cultivation with vs. without IBU (Supp. Table S4). Of these, after 1 h two secretion family proteins and one sporulation protein were significantly higher abundant in the IBU incubations as compared to the respective controls (Figure 6, Supp. Table S5). Proteins with significantly lower abundance in the IBU incubations after 1 h included a spore protein and an isocitrate lyase (Supp. Table S4). After 24 h, in total 35 proteins were significantly higher abundant in the IBU-incubated cells. These included two cytochrome P450 proteins, as well as alcohol,aldehyde, and acetyl-CoA dehydrogenases, and proteins of the amino acid metabolism (Supp. Table S5). After 24 h, in total 81 proteins were significantly lower abundant in the IBU incubations as compared to the control without IBU. These included ribosomal proteins, as well as proteins involved in sporulation, amino acid biosynthesis, and stress response proteins (Supp. Table S4).

**Figure 6:**
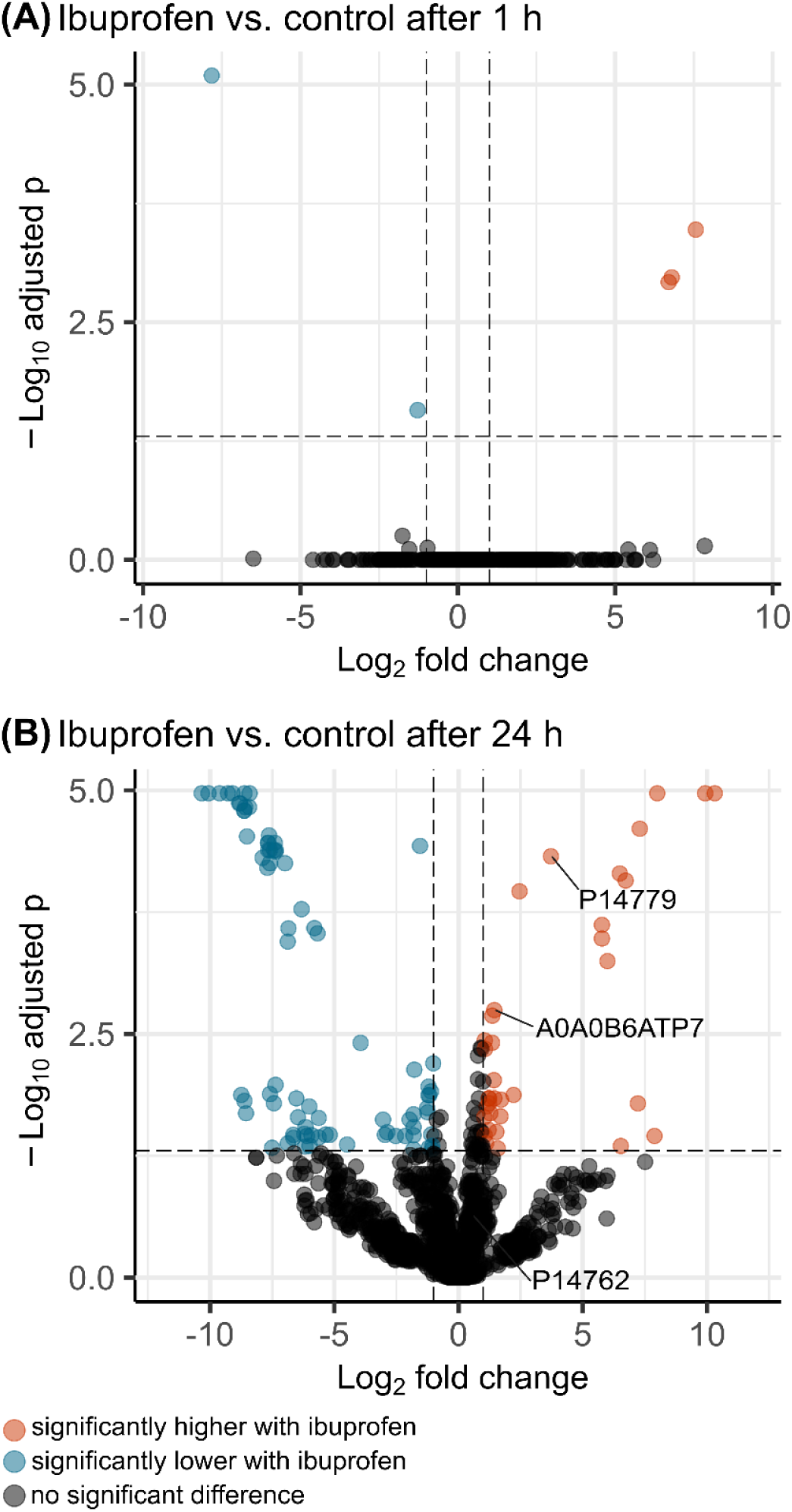
Volcano plots showing differences in the proteome of *Priestia megaterium* SBUG 518 after incubation with IBU as compared to a control without IBU. (A) Proteome comparison after 1 h incubation, (B) proteome comparison after 24 h incubation. Significance: FDR < 0.05, log_2_ fold change >1/<-1. Cytochrome P450 proteins are marked (A0A0B6ATP7: Cytochrome P450 family protein, P14779: bifunctional cytochrome P450/NADPH-P450 reductase Cyp102A1, P14762: Cytochrome P450 BM1).

## Discussion

Our experiments revealed that *P. megaterium* SBUG 518 has high potential for IBU transformation. *P. megaterium* SBUG 518 exhibited two general IBU biotransformation pathways, which we will discuss in detail below: (A) isobutyl side chain hydroxylation at two positions, and (B) conjugation with a sugar molecule. Conjugation with sugar took place almost exclusively in presence of glucose, which can be expected. More surprisingly, stationary-phase cells exhibited higher transformation rates as compared to logarithmic-phase cells.

(A) Isobutyl side chain hydroxylation yielded the products 2-OH-IBU, which accumulated in high concentrations, and CBX-IBU. While 2-OH-IBU generation by bacteria has hitherto seldomly be demonstrated (64), it is the main metabolite in urine of humans after IBU uptake (65, 66), and also a major transformation product of other higher organisms like animals (67, 68), plants (69), and fungi (70, 71). Consequently, 2-OH-IBU is widespread in the environment. It can be detected in WWTPs (72, 73), in river water and river sediments (74, 75), and in other aquatic environments (76). While CBX-IBU appears to be an unusual IBU transformation product in bacteria and fungi, it is produced by humans, some animals (65, 66, 68), and plants (69). In the environment, CBX-IBU is present in wastewater, activated sludge, and river biofilm reactors (72, 77–80). Despite that CBX-IBU is formed only after initial oxidation of IBU to 3-OH-IBU, we did not detect the primary oxidation product 3-OH-IBU, likely because the further oxidation to CBX-IBU happened too fast. Likewise, 3-OH-IBU is not, or at lower concentrations than CBX-IBU, detected in urine and environmental samples (65, 66, 68, 77, 79, 80). While we did not find additional IBU hydroxylation products in our study with *P. megaterium* SBUG 518, other organisms also produce 1-OH-IBU (*Phanerochaete chrysosporium*, Rodarte-Morales et al., 2012) and 1,2-diOH-IBU (*Trametes versicolor*, Marco-Urrea et al., 2009).

(B) Conjugation with a sugar molecule, or glycosylation, was the main driver of the high IBU transformation rate of *P. megaterium* SBUG 518 with up to 90 % transformation in 4 h. IBU was conjugated with a pyranose, yielding a glucoside. Given that this transformation took place almost exclusively in presence of glucose, the pyranose is likely to be glucose. The carboxylic group of IBU enables direct conjugate formation with the parent substrate. This is in contrast to the metabolism of many other xenobiotics, where a functional group for conjugation has to be introduced in the so-called phase I metabolism, and only then a conjugate is formed via this new part of the molecule in phase II metabolism (82). We postulate that the glycosylation of 2-OH-IBU, which we demonstrated in addition to that of IBU itself, also happens via the already-present carboxylic group, and not via the newly introduced hydroxyl group, and therefore does not adhere to “classical” xenobiotic metabolism. While IBU conjugation appears to be rare in bacteria, it is an important IBU metabolism pathway in humans, where foremost esterification via the carboxylic group of IBU and the hydroxylic group of glucuronic acid takes place, leading to acyl glucuronides (65). In addition, in human liver microsomes IBU is conjugated to glucose, and human phase I metabolites might be glycosylated, too (83). The conjugation was highly reversible: potentially, the sugar is cleaved and used as energy source by the bacteria upon depletion of external and internal energy reserves. We did not observe mineralization of IBU by *P. megaterium* SBUG 518. Mineralization by other bacteria includes ligation to CoA or acidic side chain removal and ring cleavage, as well as co-metabolic degradation, and IBU can even be used as carbon source (24, 26, 27, 30, 31, 64, 84, 85).

In summary, we propose three different pathways of IBU metabolism in *P. megaterium* SBUG 518 (Figure 7).

**Figure 7:**
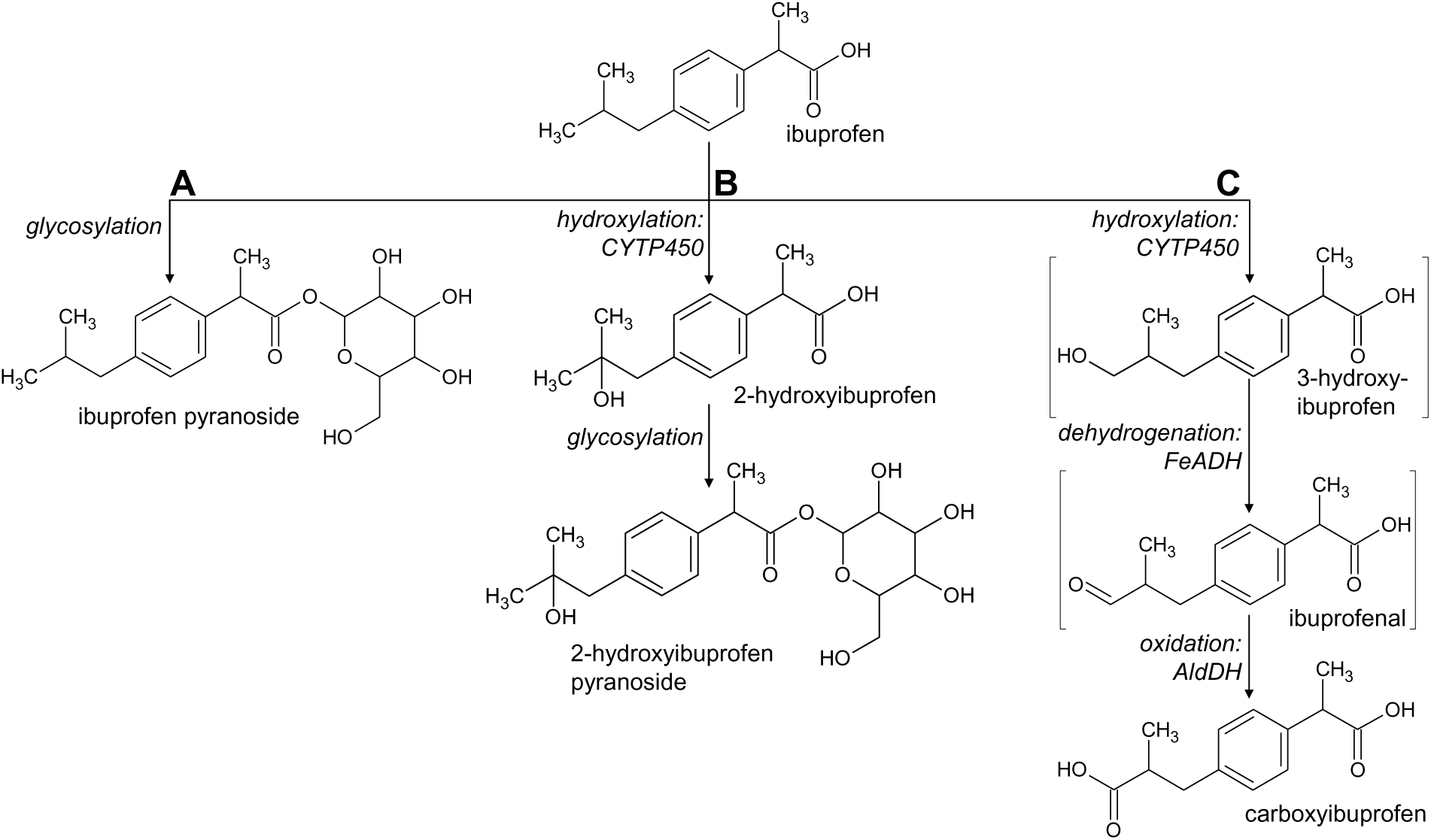
Proposed transformation pathways of IBU by *Priestia megaterium* SBUG 518 to (A) ibuprofen pyranoside, (B) 2-hydroxyibuprofen pyranoside and (C) carboxyibuprofen. Products in square brackets are proposed intermediates, which were not detected in this study. Italics: transformation reaction and proposed enzymes according to our proteomics analysis (no enzyme for glycosylation could be proposed, as transformation experiments for proteomics did not contain glucose). CYTP450: cytochrome P450, FeADH: iron-containing alcohol dehydrogenase, AldDH: aldehyde dehydrogenase.

Based on our results, conjugation of IBU likely serves as an efficient detoxification mechanism in *P. megaterium* SBUG 518, as the IBU-PYR exhibited no toxic effect in equimolar concentration to almost completely growth-abolishing IBU concentrations. Indeed, conjugates are often less biologically or chemically active than the parent compound - however, some exceptions exist, including glucuronic acid conjugates of IBU: These conjugates can directly interact with DNA and proteins, can covalently bind to plasma proteins and human albumin, and can disturb cellular functionality as far as leading to cell death (86–88). Conjugation can therefore not be regarded as universal means to decrease IBU toxicity.

Our proteomic analyses enabled us to elucidate the molecular basis for IBU transformation and IBU toxicity in *P. megaterium* SBUG 518. The two significantly higher abundant cytochrome P450 proteins in presence of IBU are likely the primary site of IBU hydroxylation in *P. megaterium* SBUG 518, as also indicated by our cytochrome P450 inhibition assay results. The bifunctional cytochrome P450/NADPH-P450 reductase Cyp102A1 (P14779, also known as P450 BM3), which we detected in over 12fold higher abundance after 24 h IBU incubation as compared to the control, is one of two different described cytochrome P450 systems in *P. megaterium* (89, 90). It consists of a P450 fatty acid hydroxylase and a mammal-like, albeit soluble, diflavin NADPH-P450 reductase in a single enzyme (91). P450 BM3 can be induced by IBU (92), and this enzyme can transform a variety of drugs to different metabolites (93). For example, in *P. megaterium* ATCC 14581, P450 BM3 oxidizes fatty acids, long-chain alcohols, and amides (94, 95).

Also, significantly higher abundant in IBU incubations were iron-containing alcohol dehydrogenases (FeADHs), which might generate ibuprofenal from 3-OH-IBU. Ibuprofenal could then be oxidized further to CBX-IBU via an aldehyde dehydrogenase. In *Clostridium*, which is closely related to *Bacillus*, FeADHs can have catalytic functions as propanediol dehydrogenase, butanol dehydrogenase or 4-hydroxybuyrate dehydrogenase (96–98). Therefore, these enzymes utilize substrates similar to the isobutyl and propanoic acid residues of IBU, supporting our functional hypothesis.

The two up-regulated lipid/propionate metabolism enzymes, 3-hydroxypropionyl-coenzym A dehydratase and enoyl-CoA hydratase/isomerase, could participate in transformation of propionate to acetyl-CoA (via acrylate and 3-OH-propionate, as alternative to the methylmalonyl-CoA-pathway to succinyl-CoA), which could then be further metabolized via acetyl-CoA dehydrogenase. This is the case in several bacteria and eukaryotic organisms (99, 100). Propionate can be split off from side chains of IBU by several bacteria (as shown e.g. in this study by the formation of 4-carboxy-α-methylbenzeneacetic acid by *M. neoaurum* SBUG 109). While in *P. megaterium* SBUG 518 we hitherto did not find C3-dealkylated products of IBU, at least four of the formed transformation products have not been identified yet. Alternatively, it cannot be ruled out that some of these enzymes could also directly transform the IBU side-chains without any off-splitting of C3-units or were induced during degradation of valine or isoleucine also resulting in propionyl-CoA formation. In the IBU-degrading, Gram-positive *Patulibacter* I11, proteins involved in fatty acid metabolism were also suggested to be involved in IBU degradation (24).

In addition, we detected two HlyD family transporters only after incubation of *P. megaterium* SBUG 518 with IBU. These transporters likely transport IBU and its transformation products out of the cells. HlyD family transporters belong to ABC transporters, which in Gram-positive bacteria are often used to expel xenobiotics (101).

Coupled to this direct response to IBU, we also found proteome profile changes suggestive of a more general physiological response of *P. megaterium* SBUG 518 to IBU presence, i.e., a pleiotropic effect of the drug. Single stranded DNA specific exonuclease, which was after 24 h only detected in IBU-containing incubations, but not in the respective controls, suggests that DNA damage occurred in IBU presence, necessitating DNA repair. In *Escherichia coli*, several NSAIDs impact DNA replication and repair by inhibition of DNA polymerase III beta subunit (102). It remains to be elucidated whether IBU has a similar mode of action in *P. megaterium* SBUG 518. Interestingly, the relaxosome subunit MobC was detected in 10fold higher concentration after 24h incubation with IBU as compared to the control, suggesting that horizontal gene transfer is promoted by IBU presence.

Moreover, IBU seems to impact spore formation in *P. megaterium* SBUG 518: Several spore-forming proteins were significantly lower abundant after 24 h incubation with IBU as compared to the controls. Corroborating this, we noticed that logarithmic-phase cells did not sporulate when incubated with IBU without glucose (data not shown). While data on the impact of IBU on cell wall turnover and sporulation seems to be lacking, several studies describe negative impacts of this drug on the membrane integrity of pro- and eukaryotic microbial species (103–105). The high resistance of *Bacillus thuringiensis* B1(2015b) against IBU, on the other hand, is likely based on changes in the membrane composition (85). In a broader context, sporulation in *B. subtilis* is inhibited by unsaturated fatty acids (106), which bear structural and metabolic resemblance to IBU.

IBU also seems to interfere with the amino acid metabolism and protein synthesis of *P. megaterium* SBUG 518, as several proteins involved in amino acid metabolism were significantly lower or higher abundant, and ribosomal proteins lower abundant after incubation with IBU as compared to the controls. Potential reasons for this are shifts in protein expression patterns, due to the synthesis of detoxifying and transporting proteins as well as repair proteins, and potential interference of IBU with gene expression. At the same time, in *P. megaterium* SBUG 518, the fatty acid metabolism is likely negatively impacted by IBU, as we detected the fatty acid biosynthesis enzymes malonyl CoA-acyl carrier protein transacylase, enoyl-[acyl-carrier-protein] reductase in significantly lower abundance after IBU incubation. While it remains unclear whether this is due to general energy shortage or due to direct effects of IBU in *P. megaterium* SBUG 518, also in oral pathogenic bacteria IBU potentially interacts with proteins involved in fatty acid metabolism (107).

Interestingly, the oxidative stress proteins catalase and alkyl hydroperoxide reductase were significantly lower abundant in *P. megaterium* SBUG 518 after 24 h IBU incubation. IBU has been shown to induce catalase activity in *P. megaterium* (108), and cytochrome P450 activity leads to oxygen radical formation (109), which makes the lower abundance of oxygen radical detoxifying enzymes in *P. megaterium* SBUG 518 in IBU presence unexpected. However, reactive oxygen species themselves might be involved in cytochrome P450 BM3 induction, as induction in a *P. megaterium* strain decreased when cells were incubated with IBU and external catalase (108). Therefore, protein expression regulation might be more complex than anticipated. Additionally, an antioxidant effect of IBU itself has been described (110), which might add to this protein abundance pattern. Moreover, IBU itself might interfere with the respective gene expression, actually exacerbating oxidative stress in *P. megaterium* SBUG 518.

Taken together, while *P. megaterium* SBUG 518 efficiently transforms IBU as a means of detoxification, the drug still elicits various and apparently detrimental effects on the physiology of this bacterium.

## Conclusion

*P. megaterium* SBUG 518 exhibits IBU transformation and detoxification mechanisms similar to those in humans. Therefore, *P. megaterium* SBUG 518 seems to be well suited as a model organism for further research of drug metabolism, not least because its cytochrome P450 monooxygenase highly resembles the corresponding eukaryotic enzymes. The reversibility of the fast glycosylation of IBU can have practical consequences: It is an open question why, despite the apparently nearly complete removal of IBU in WWTPs, it can still be detected in various surface waters. A temporary “masking” of IBU as IBU-PYR could explain this. Therefore, kinetic data of IBU transformation should take into account possible conjugate formation. Future studies on IBU as antimicrobial agent should shed more light on the mode of action of this drug in different microbial organisms.

## Acknowledgements

The authors are grateful to Sabryna Junker, Veronika Hahn, Anne Reinhard and Stefan Bock for excellent technical assistance.

## Supplementary Material

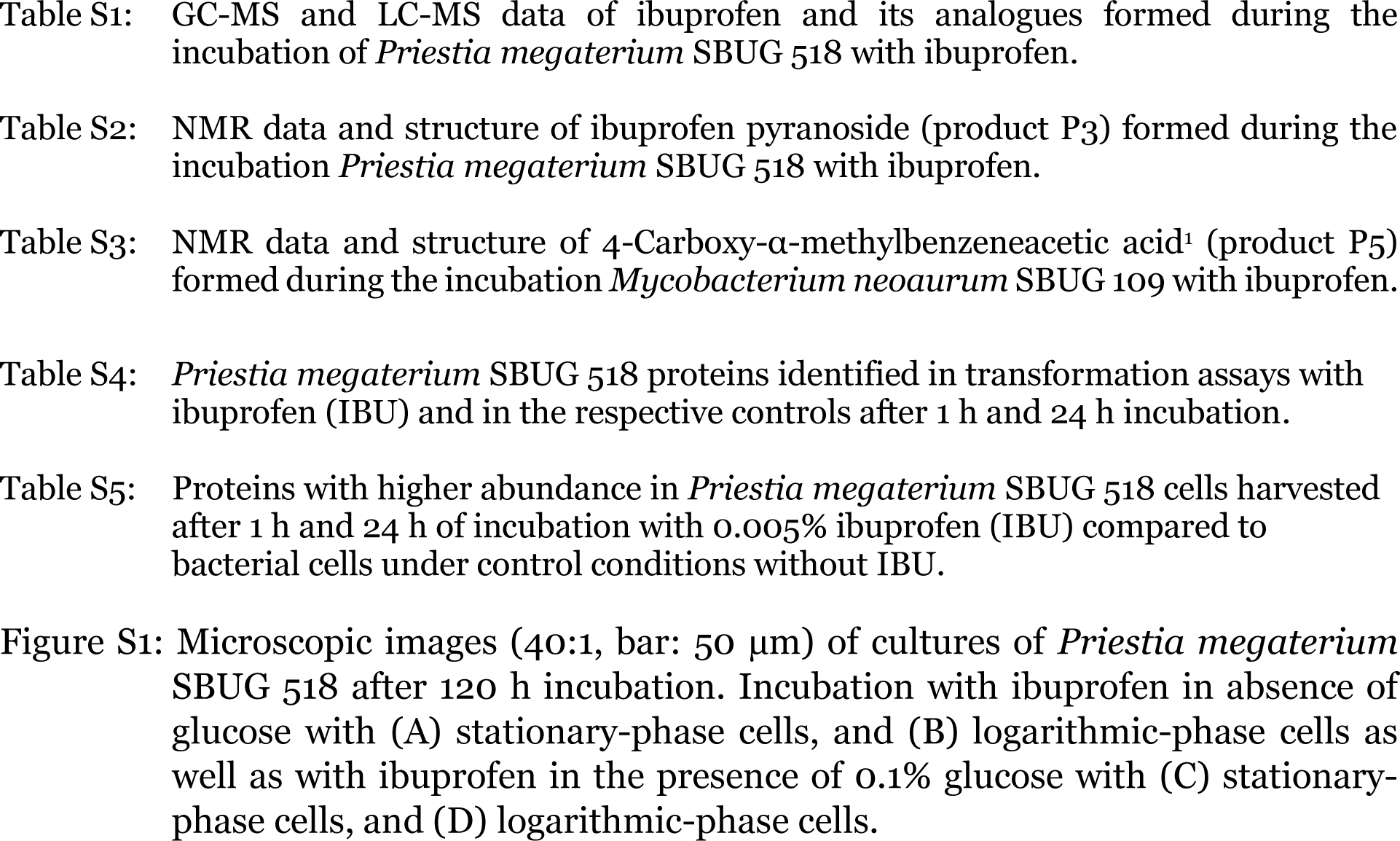

